# The c-di-GMP effector FleQ controls alginate production by repressing transcription of the *algD* gene in *Azotobacter vinelandii*

**DOI:** 10.1101/2024.09.30.615961

**Authors:** Víctor V. Barrios-Rafael, Carlos L. Ahumada-Manuel, Scherezada Orgaz-Hidalgo, Jessica Nava-Galeana, Josefina Guzmán, Soledad Moreno, Victor Bustamente, Cinthia Núñez

**Affiliations:** Departamento de Microbiología Molecular, Instituto de Biotecnología, Universidad Nacional Autónoma de México (UNAM). Cuernavaca, Morelos, México

## Abstract

*Azotobacter vinelandii*, belonging to the *Psedomonadaceae* family, produces the exopolysaccharide alginate during its vegetative growth and also during its differentiation process to form desiccation resistant cysts. The second messenger c-di-GMP regulates a vast array of cellular processes. It is produced by diguanylate cyclases (DGC) and degraded by phosphodiesterases (PDE). In *A. vinelandii* the absence of the AvgReg DGC impairs alginate synthesis while the absence of the PDE MucG increased alginate production. The effect of the different c-di-GMP levels was solely attributed to its essential role in activating the alginate polymerase complex. Here in we investigated the possible role of c-di-GMP in the control of *algD* transcription, encoding the key enzyme of the alginate biosynthetic pathway. At artificially high or reduced levels of c-di-GMP *algD* transcription was increased, or almost abrogated, respectively. Therefore, the role of the transcriptional regulator FleQ, one of the best characterized c-di-GMP effectors, was investigated. Alginate production increased in the Δ*fleQ* relative to the wt strain, which agreed with increased *algD* transcription. These phenotypes were rescued in the Δ*fleQ*/*fleQ*+ complemented strain indicating a FleQ repressing effect. EMSA assay showed that FleQ was able to directly bind to the regulatory region of *algD* which agreed with the presence of a FleQ binding site overlapping its RpoS-dependent promoter. In *A. vinelandii* the c-di-GMP is also necessary for expression of alginate C-5 epimerase genes, essential for structuring mature cysts. Out data revealed that FleQ is not an intermediary in this regulation since its absence did not impair mature cyst formation nor affected expression of *algE1-6* genes. Collectively, our results support a model in which FleQ is the intermediary in the regulation of *algD* by c-di-GMP, exerting a direct repressing effect and reveal the existence of a FleQ-independent regulatory mechanism for the control of *A. vinelandii* encystment.

## Introduction

*Azotobacter vinelandii,* belonging to the *Pseudomonadaceae* family is a nitrogen-fixing bacterium that is motile by the presence of peritrichous flagella. The *Azotobacter* genus undergoes a differentiation process for the formation of desiccation-resistant cysts [1]. Upon encystment, biochemical, metabolic, and morphological changes are observed, including the loss of the flagella and the arrest of nitrogen fixation, among many others. The central body of the mature cyst is surrounded by two protective layers, the exine and the intine, containing a high proportion of the exopolyssacharide alginate and essential for dehydration resistance [2].

The alginate is a linear poly-saccharide composed of two monomers, the α-L-mannuronate (M) and its C-5 epimer the β-D-guluronate (G). Alginate synthesis is started from Fructose-6-P that after three enzymatic steps in the cytosol, is transformed to the activated precursor GDP-mannuronate, that serves as the substrate for the alginate polymerase complex located in the inner membrane [3–5]. The product of the *algD* gene catalyzes the oxidation of GDP-mannose to GDP-mannuronate. This reaction is the rate limiting step of the pathway and hence, expression of *algD* is subjected to a complex genetic regulation [3]. Transcription of *algD* starts from three promoters recognized by the sigma factors RpoS (for promoter P1) or AlgU (for promoter P2) [6–8]; the nature of the third signal detected as a putative transcriptional start site is currently unknown [9]. In contrast to *P. aeruginosa*, where several transcriptional regulators have been described for *algD*, such as AmrZ, AlgB or AlgR [10–13], no transcriptional regulators have been so far reported for *A. vinelandii*, that directly recognize the regulatory region of *algD* [3].

Previous work from our laboratory revealed that the second messenger bis (3’,5’)-cyclic dimeric guanosine monophosphate (c-di-GMP) exerts a positive effect on the amount of alginate produced by *A. vinelandii* and on its molecular mass [14, 15]. The second messenger c-di-GMP controls a vast array of cellular processes in bacteria, including the transitions from the motile to the sessile style, biofilm formation or the progression of the cell cycle [16]. It is produced from GTP by the activity of diguanylate cyclases (DGC) and degraded by phosphodiesterases (PDE). C-di-GMP effectors convert changes in c-di-GMP concentration into cellular responses and such effectors include transcription factors, riboswitches or signaling proteins [17]. Under the vegetative growth of *A. vinelandii*, a c-di-GMP module composed by the DGC AvgReg and the PDE MucG provides the c-di-GMP necessary for alginate production [14]. The absence of MucG increased the levels of c-di-GMP and those of alginate production of higher molecular mass, when compared to the wild type strain [14, 18]. Since the polymerase complex Alg8-44 is activated by binding of c-di-GMP to the co-polymerase Alg44 [19, 20], these alginate phenotypes were attributed to increased activation of the polymerase complex. Higher *algD* expression was observed in the absence of the PDE MucG [18]. However, the possible regulatory effect of c-di-GMP upon *algD* transcription had not been addressed.

In *A. vinelandii* the c-di-GMP was also found to be essential for the formation of mature cysts. The alginate protective layers of the cysts, essential for desiccation resistance, contain different proportions and distributions of G residues which result from the activity of a family of C-5 epimerases, encoded by the genes *algE1-6*. The second messenger c-di-GMP was necessary for expression of *algE1-6* genes and, therefore, to produce mature cysts resistant to desiccation [15].

FleQ is one of the best c-di-GMP effectors so far characterized in *Pseudomonas* species [21]. Although FleQ shares significant homology to members of the NtrC family, containing an N-terminal receiver domain (REC), a central ATPase (AAA+) domain, and a C-terminal HTH DNA binding motif [22], its activity responds to changes in the levels of c-di-GMP altering target promoter activity and it is also influenced by the FleN antagonist and ATP hydrolysis [23–25]. In *P. aeruginosa* FleQ is the master regulator of flagellar motility and exopolyssacharide production, mediating the transition from the planktonic to the biofilm lifestyles. At low c-di-GMP, FleQ activates flagellar genes but represses genes encoding biofilm matrix components, such as the Pel polysaccharide. However, at high c-di-GMP, FleQ is unable to active the flagellar genes but upregulates expression of genes encoding matrix components necessary for biofilm formation [21].

In this study the role of FleQ as a possible intermediary in the regulation of *algD* transcription and mature cyst formation by c-di-GMP in *A. vinelandii* was investigated. A Δ*fleQ* mutant showed three times higher levels of alginate because of a direct repression of *algD* transcription. In contrast, the Δ*fleQ* mutant was able to form mature cysts resistant to desiccation, implying that a different regulatory mechanism responding to c-di-GMP controls transcription of *algE1-6* genes and cellular differentiation in this bacterium.

## Materials and Methods

### Strains and cultivation conditions

The wild-type strain AEIV of *A. vinelandii* was used [26] and was routinely cultivated in Burk’s medium; sucrose (20 gL-1) was used as the carbon source (Burk’s-sucrose medium). The composition of the growth medium has been reported previously [27]. Cultures were incubated at 30 oC and 200 rpm.

Resistance to desiccation of *A. vinelandii* cysts was determined as reported previously [6, 15]. *A. vinelandii* transformation was conducted following a protocol reported previously [28, 29] with some modifications detailed elsewhere [18]. The final concentration of antibiotics for the selection of *A. vinelandii* transformants on Burk’s-sucrose plates was gentamacyn (Gm) 1 μg mL-1; tetracyclin (Tc) 30 μg mL-1; kanamycin (Km), 1 μg mL-1.

### Standard Techniques

Genomic DNA isolation was conducted with the Bacterial DNA preparation kit following instructions of the manufacturer (Jena Bioscience). The high-fidelity Phusion DNA polymerase (Thermo Fischer Scientific) was used for all PCR reactions and were confirmed by DNA sequencing. DNA sequencing was carried out with fluorescent dideoxy terminators using a cycle sequencing method and a 3500xl analyzer (Applied Biosystems).

### Analytical methods

Protein concentration was determined by the Lowry method [30]. The activity of β-galactosidase in *A. vinelandii* cells was determined as reported before [18]. Alginate concentration was determined as described [14], using the spectrophotometric determination of uronic acids with carbazole [31]. All experiments were conducted in triplicates; the results presented are the averages of the independent runs. Statistical analysis was carried out using a Student’s t-test (p = 0.05).

### Construction of mutants

The *fleQ* mutant was constructed by PCR amplification of a DNA fragment of 2362 bp containing the *fleQ* gene, using chromosomal DNA from strain AEIV as template and the primer pairs FleQ-F (5’-GGC TTG GCT GGA AGC ATT G-3’)/FleQ-R (5’-GGC AGT CAG CAC ACG ATA G-3’). The PCR product was subsequently cloned into vector pJET1.2/Blunt (Thermo Fischer), rendering plasmid pLA62. A SphI-EcoRV *fleQ* internal fragment of 1229 bp from plasmid pLA62, was replaced by a Gm cassette, released with SmaI endonuclease from plasmid pBSL190 [32]. Before cloning the resistance cassettes, the pLA62 plasmid, excised with SphI-EcoRV, was previously made blunt using the Klenow enzyme. The resultant plasmid was named pLA498 (Δ*fleQ*::Gm). The orientation of the resistance cassettes in plasmid pLA498 is the same as that of *fleQ*, preventing polar effects of downstream genes. Plasmid pLA498 previously linearized with ScaI endonuclease, was used to transform competent AEIV cells. The resultant Δ*fleQ* mutant was named CLAM522 (Δ*fleQ*::Gm).

Genetic complementation of the Δ*fleQ*::Gm mutant was conducted by integration a wild type copy of *fleQ* into the genome of CLAM522. For this purpose, the DNA product of 2362 bp containing the *fleQ* gene and its regulatory region, was cloned into vector pUMATc5’-3’ (Tc) previously excised with SmaI [33], rendering plasmid pUMA-fleQ. This plasmid, previously linearized with ScaI, was introduced by transformation into CLAM522. After a double homologous recombination, the *fleQ* gene was integrated into the *A. vinelandii* genome within a neutral locus (*melA*). The complemented strain (Δ*fleQ*/*fleQ+)* was named SM654.

The transcriptional *algD-lacZ* fusion present in strain A2 [34] was used to assess the influence of intracellular levels of c-di-GMP or that of FleQ on *algD* transcription. Competent cells of A2 strain were transformed with chromosomal DNA from strains Δ*fleQ* (Gm^r^), DGC+ (Tc^r^) or PDE+ (Tc^r^), followed by the selection of transformants in the presence of the corresponding antibiotic. The resulting strains were named JG621, JG623 and JG628, respectively.

### Real-Time qPCR

The relative mRNA levels of the *algD* gene or *algE1* to *algE6* (*algE1-6*) genes, were determined by RT-qPCR following a protocol described previously [15], using total RNA extracted from cells cultivated in Burk’s-sucrose (vegetative conditions) or Burk’s-n-butanol medium (encysting conditions) for 24 h. Total RNA was extracted as reported [35]. The primer pairs used were algD-RT3-F (5’-TTCGGACTGGGCTATGTAGG-3’)/algD-RT3-R (5’GCCCTGATTGATCATGTCG-3’) for *algD* and FwRT-algE1-6 (5’-CACGAGCAGACCATCAACCTG-3’)/ RvRT-algE1-6 (5’-ATGTTGAAGCCGTGGCGGTCGTTG-3’), for genes *algE1-6*. The relative levels of *algD* and *algE1-6* were determined by comparing the amount of each mRNA using *gyrA* (Avin_15810) mRNA as an internal control. Three biological replicates (independent cultures) were conducted, with three technical replicates for each one. The quantification technique employed to analyze the generated data was the 2-DDCT method [36].

### Identification of potential FleQ binding sites

13 sequences identified as FleQ binding sites in *P. aeruginosa* [37] were used to build the position-specific scoring matrix and sequence *logo* of FleQ using MEME (Multiple Em for Motif Elicitation), one occurrence per sequence was searched [38]. The position-specific scoring matrix was used as input to FIMO (Find Individual Motif Occurrences) algorithm for searching FleQ binding sites in the 5’ UTR gene sequences of *A. vinelandii* DJ strain [39]. The 5’ UTR gene sequences of *A. vinelandii* were obtained using bedtools getfasta command from bedtools toolset [40], encompassing 400 and 50 nucleotides upstream and downstream of the genomic start coordinate of each gene found in the *A. vinelandii* DJ annotation file, respectively.

### Expression and Purification of the FleQ-His protein

FleQ was purified as a recombinant 6XHis tag at the N terminus (FleQ-His). The *fleQ* gene was PCR amplified using the primer pairs fleQ-F-NdeI (5’-CATATGATGTGGCGTGACATAAAAATCCTCC-3’)/fleQ-R-BamHI (5’-GGATCCCGCAGCGAGGTTTAACAC-3’) and cloned into pET-28a vector previously excised with NdeI and BamHI endonucleases, leaving an N-terminal His6 tag. The resulting plasmid was named pET-FleQ. BL21 (DE3) *E. coli* strain (Invitrogen), carrying pET-FleQ was grown in LB Km at 37°C until an OD_600_ of 0.6. At this point, 0.05 mM of IPTG (isopropyl-D-thiogalactopyranoside) was added to induce expression of FleQ-His and was incubated 15°C for 16 h. Cells were harvested by centrifugation at 4000 xg for 10 min, washed once with cold lysis buffer (50 mM NaH_2_PO_4_, 300 mM NaCl, 10 mM imidazole [pH 8]; pH 8) and disrupted by sonication in the same buffer. The supernatant was recovered by centrifugation at 4000 xg for 30 min at 4°C. The crude extract was loaded to a column containing the nickel resin Ni-nitriloacetic acid agarose previously equilibrated with lysis buffer. The column was washed ten times with washing buffer (50 mM NaH_2_PO_4_, 300 mM NaCl, 20 mM imidazole [pH 8]; pH 8) and the protein was eluted with elution buffer I (50 mM NaH_2_PO_4_, 300 mM NaCl, 250 mM imidazole [pH 8]; pH 8). The protein was concentrated by using Amicon Ultra 50K device (Merck Millipore) and stored in elution Buffer II (20 mM Tris-HCl [pH 8], 300 mM NaCl, 20% glycerol). Protein concentration was determined by the Bradford technique using bovine serum albumin as standard. SDS-PAGE revealed the purified FleQ-His protein.

### Electrophoretic mobility shift assay (EMSA)

EMSA assay was performed as previously described, using the non-radioactive method [41, 42]. A DNA fragment of the regulatory region of *algD*, containing the RpoS-dependent promoter, including the potential FleQ binding site, was PCR amplified using the primer pairs algD-EMSA-F1.21 (5’-TACGGCAATCCCATTGCTG -3’)/palgD-R (5’-CCGAAAATGCTGATACGC-3’) and chromosomal DNA of strain AEIV. Binding reactions were performed by mixing 100 ng of the DNA fragment with increasing concentrations of purified FleQ-His protein in binding buffer (10 mM Tris [pH 8], 8 mM magnesium-acetate tetrahydrate, 50 mM KCl, 10 µg/mL bovine serum albumin and 5% glycerol), in a total volume of 20 mL. Binding reactions were incubated at room temperature for 20 min before loaded onto a 6% nondenaturing acrylamide gels at 85V with 0.5X Tris-borate-EDTA buffer. As a negative control the regulatory region of the *Salmonella* Typhimurium *eutR* gene was included. This fragment of 400 bp, was PCR amplified using the primer pairs F2eutR-EcoRI (5’-CTTGAATTCGTTTGCTCAGTCATCAAGTGC-3’)/R2eutR-BamHI (5’-CTTGGATCCCGCTGATGAACATTGTCCACC-3’). Migration of the DNA bands was visualized by staining with EtBr under UV light.

### Detection of AlgE C-5 epimerases

AlgE epimerases associated to the surface of differentiated cells were detected by Western Blot, using anti-AlgE4 antibodies and following a protocol reported previously [43]. Protein extracts from the surface of differentiated cells were prepared as described [15].

## Results

### The absence of FleQ favors alginate production

The *A. vinelandii* genome encodes a FleQ protein of 501 amino acids showing 71% identity when compared to that of *P. aeruginosa*. The motifs important for the binding of dimeric c-di-GMP, interaction with RpoN, or the active sites for the AAA+ domain are well conserved between the two proteins (S1 Fig). Similarly, sequence of the FleQ HTH domains and their structure prediction by AlphaFold are conserved (S2 Fig). It is of interest to note that a 21 amino acid motif is present at the C-terminus of FleQ proteins from the *Azotobacter* but not from the *Psedomonas* genus (S2 Fig). However, the functional relevance of this feature is not known.

To investigate the role of FleQ in *A. vinelandi,* a Δ*fleQ* deletion mutant was constructed. A hyper mucoid colony phenotype was readily apparent in the Δ*fleQ* mutant suggesting a negative effect of FleQ on alginate production (Fig 1A). Indeed, alginate production by the Δ*fleQ* mutant cultured in Burk’s-sucrose liquid media increased three-fold relative to wt strain (Fig 1B). A complemented Δ*fleQ* derivative, carrying a wt copy of *fleQ* (Δ*fleQ* /*fleQ*^+^), under its own promoter, was constructed and this strain showed wild type levels of alginate, thus confirming the negative role of FleQ in this biosynthetic pathway (Fig 1A). Cell growth was slightly decreased in the Δ*fleQ* mutant with respect to wt strain likely because of the over production of alginate, but it was restored in the Δ*fleQ*/*fleQ^+^* complemented strain (Fig 1C).

**Fig 1.**
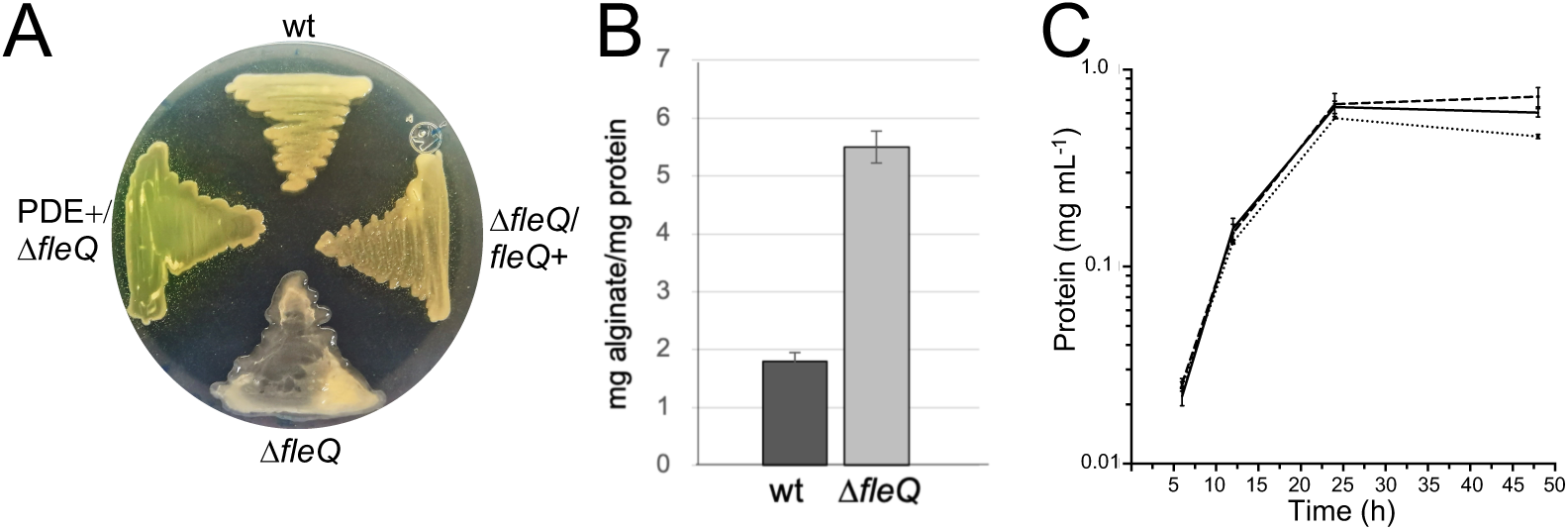
FleQ exerts a negative influence on alginate production. A. Growth of the wild type strain (wt), the Δ*fleQ* mutant and its derivatives Δ*fleQ*/*fleQ*+ and PDE+/ Δ*fleQ* on Burk’s-sucrose plates for 48 h. The hyper mucoid colony phenotype of strain Δ*fleQ* is readily apparent. B. Quantification of the alginate produced by wt strain and the Δ*fleQ* mutant grown in Burk’s-sucrose liquid medium for 48 h. C. Growth kinetics of wt strain (solid line), the Δ*fleQ* mutant (dotted line) and of its complemented strain Δ*fleQ/fleQ+* (dashed line) in Burk’s-sucrose medium.

### FleQ inhibits *algD* transcription

Given the negative effect of FleQ on alginate production, its influence on *algD* transcription was investigated. A transcriptional *algD-lacZ* fusion was used to evaluate the effect of FleQ along the growth curve. As can be seen in Fig 2A, a consistent higher level of β-galactosidase was detected in the absence of FleQ, particularly during the logarithmic phase of growth (12 and 24 h). This result was further supported by qRT-PCR using mRNA extracted from strains wt and Δ*fleQ* cultured in liquid Burk’s-sucrose medium for 24 hr. The relative mRNA levels were 1.8-fold higher in the Δ*fleQ* mutant (Fig 2B), indicating a negative effect of FleQ on *algD* transcription.

**Fig. 2.**
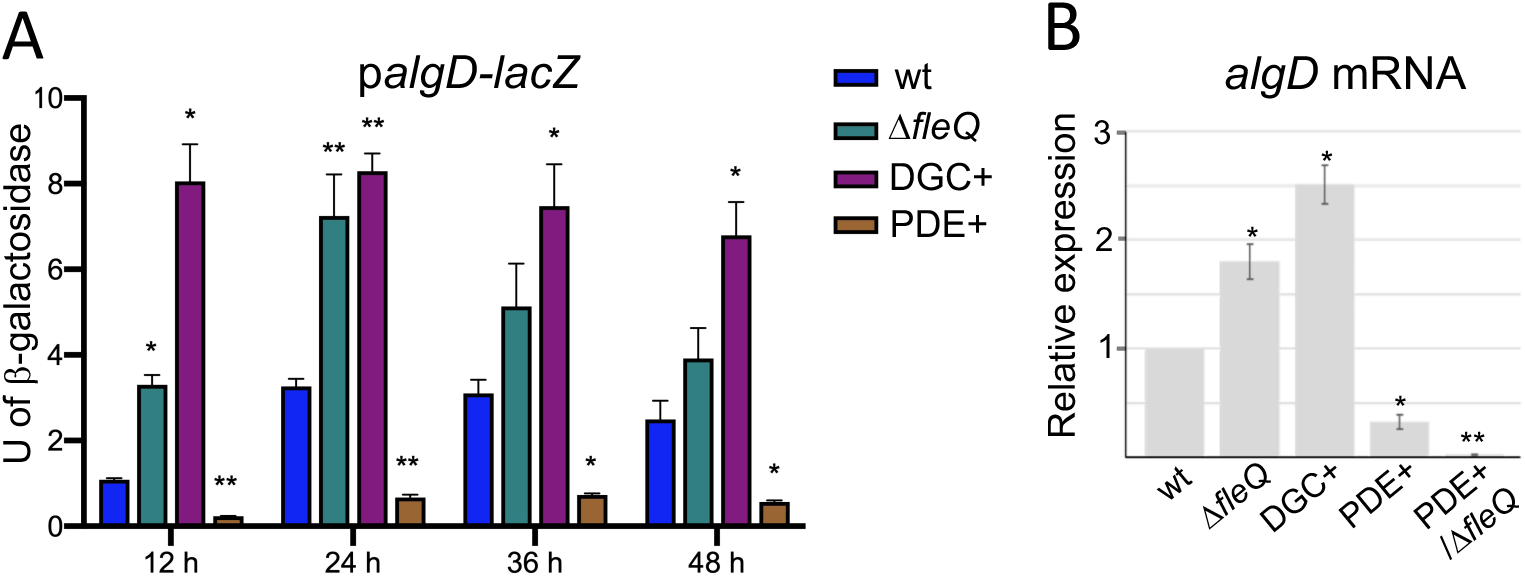
*algD* transcription is under the control of FleQ and responds to the levels of c-di-GMP. A. Kinetics of *algD* transcription using an *algD-lacZ* transcriptional fusion in strains wt and Δ*fleQ*, or in strains showing high (DGC+) or reduced (PDE+) levels of c-di-GMP. B. *algD* mRNA quantification by RT-qPCR using RNA extracted from strains wt, Δ*fleQ*, DGC+, PDE+ and PDE+/Δ*fleQ*.

### *algD* transcription is modulated by the levels of c-di-GMP

In agreement with the negative influence of FleQ on *algD* transcription, we found that expression of *algD* was positively modulated by the levels of c-di-GMP. Transcription of *algD* along the growth curve, using the transcriptional *algD-lacZ* fusion, correlated with artificially high or reduced levels of c-di-GMP obtained from over-expression of the DGC AvGREg (DGC+) or the PDE Avin_50640 (PDE+), respectively (Fig 2A) [14]. It is worth noting that *algD* transcription was almost abrogated in strain PDE+. This data was further supported by qRT-PCR analysis, using total RNA extracted from strains DGC+ and PDE+ cultivated in Burk’s-sucrose medium for 24 h. Transcripts of *algD* were 2.5-fold higher and 3-fold reduced in the DGC+ and PDE+ strains, respectively relative to the wt background (Fig 2B), revealing the positive effect of c-di-GMP on *algD* and contributing to explain the alginate over-producing phenotype previously reported by strain DGC+ or the reduced alginate levels in strain PDE+ [14].

### FleQ is a direct repressor of *algD* transcription

A MEME/MAST analysis was conducted as described in Materials and methods, aimed at identifying potential targets of FleQ in the *A. vinelandii* genome (S1 Table). Given the conservation of the HTH domain between the FleQ proteins from *A. vinelandii* and *P. aeruginosa* (S1 and S2 Fig), this analysis was carried out using *P. aeruginosa* experimentally demonstrated FleQ binding sites. A total of 226 sites in the *A. vinelandii* genome were identified with a p-val <0.0001, among them a potential binding site in the regulatory region of *algD* (p-val 4.30E-05), overlapping its RpoS promoter (Fig 3A). To confirm this prediction, an EMSA assay was conducted using the purified *A. vinelandii* FleQ protein. In the presence of FleQ, the migration of a DNA fragment of the regulatory region of *algD* was retarded indicating the formation of a DNA/protein complex (Fig 3B). The binding of FleQ seems to be specific as FleQ did not retard a DNA fragment used as negative control. In addition, the regulatory region of the *P. aeruginosa pelA* gene was included as a positive control (S3 Fig). We found that the *A. vinelandii* FleQ protein was able to recognize the *pelA* FleQ binding sites, thus implying that FleQ from *A. vinelandii* and *P. aeruginosa* are able of recognizing similar consensus sequences.

**Fig. 3.**
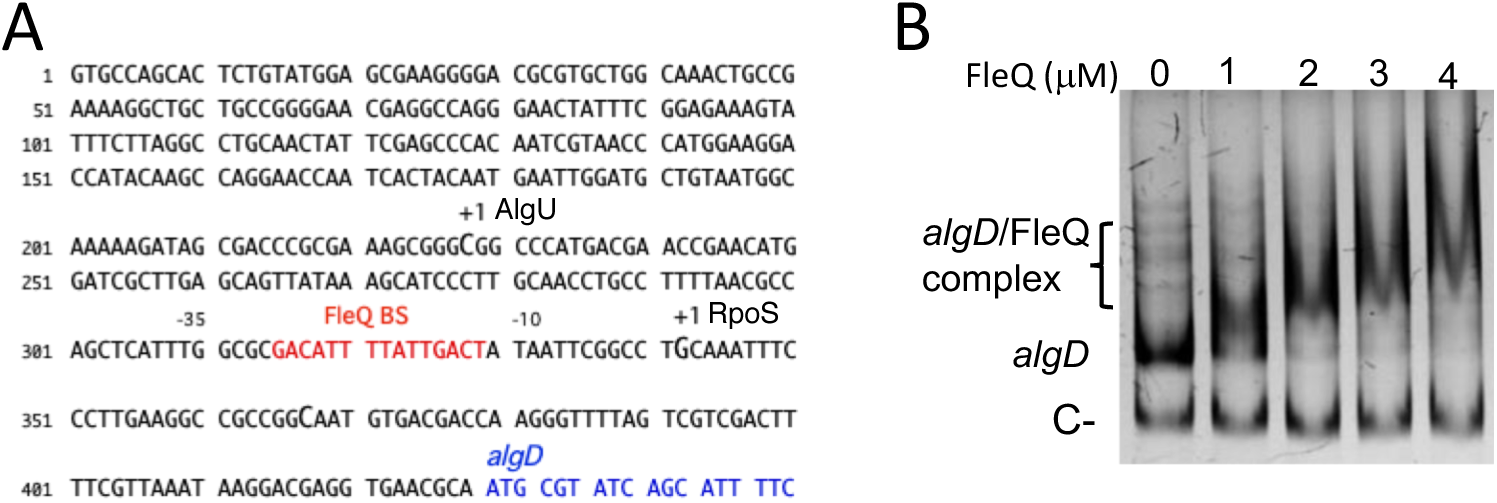
FleQ binds to the *algD* regulatory region. A. DNA sequence of the *algD* regulatory region. The transcription start sites (+1) previously identified for promoters RpoS and AlgU are indicated along with the putative FleQ binding site, overlapping the RpoS promoter. The structural region of *algD* is in blue. B. Non-radioactive EMSA assay to detect the binding of FleQ to the regulatory region of *algD*. The DNA fragment was incubated with increasing concentrations of FleQ. As a negative control (C-), a fragment of the *Salmonella Typhimurium eutR* gene was included. The migration of the DNA fragments was revealed by staining with ethidium bromide.

### c-di-GMP and the control of *algD*

Our data revealed that FleQ inhibits alginate production by directly repressing *algD* transcription. In addition, c-di-GMP favored expression of *algD* likely because of relieving the repressing effect exerted by FleQ. In the PDE+ strain, low levels of c-di-GMP reduced *algD* expression by tree-fold (Fig 2B). Therefore, we wonder if expression of *algD* could be restored under low levels of c-di-GMP but in the absence of FleQ. To this end, a Δ*fleQ* mutant was constructed in the PDE+ genetic background. However, neither alginate production (Fig 1A) or *algD* expression (Fig 2B) were reestablished in this strain implying that the second messenger c-di-GMP controls *algD* expression by an additional unknown mechanism.

### Expression of *algE1-6* genes responds to the c-di-GMP levels

In *A. vinelandii*, the DGC MucR was found essential for the transcription of the *algE1-6* genes, encoding alginate C-5 epimerases [15]. However, a direct link between c-di-GMP and *algE1-6* expression had not been established. To address this question, detection by Western Blot of the AlgE1-6 proteins on the surface of differentiated cells was conducted in the genetic backgrounds of DGC+ or PDE+ strains. The higher levels of c-di-GMP in strain DGC+ did not change the detection pattern of the epimerases attached to the cyst surface (Fig 4A); in contrast, reduced levels of c-di-GMP abolished the detection of these enzymes in strain PDE+. This effect seems to be at the transcriptional level. Transcripts of *algE1-6* genes were measured as described in Materials and Methods, using a single pair of primers previously reported [44], and able to anneal to *algE1-6* genes due to the modularity of the encoded proteins [45]. *algE1-6* transcripts were almost undetectable in strain PDE+ (Fig 4B) but two-fold higher in strain DGC+, confirming the positive role of c-di-GMP on the expression of *algE1-6* genes.

**Fig 4.**
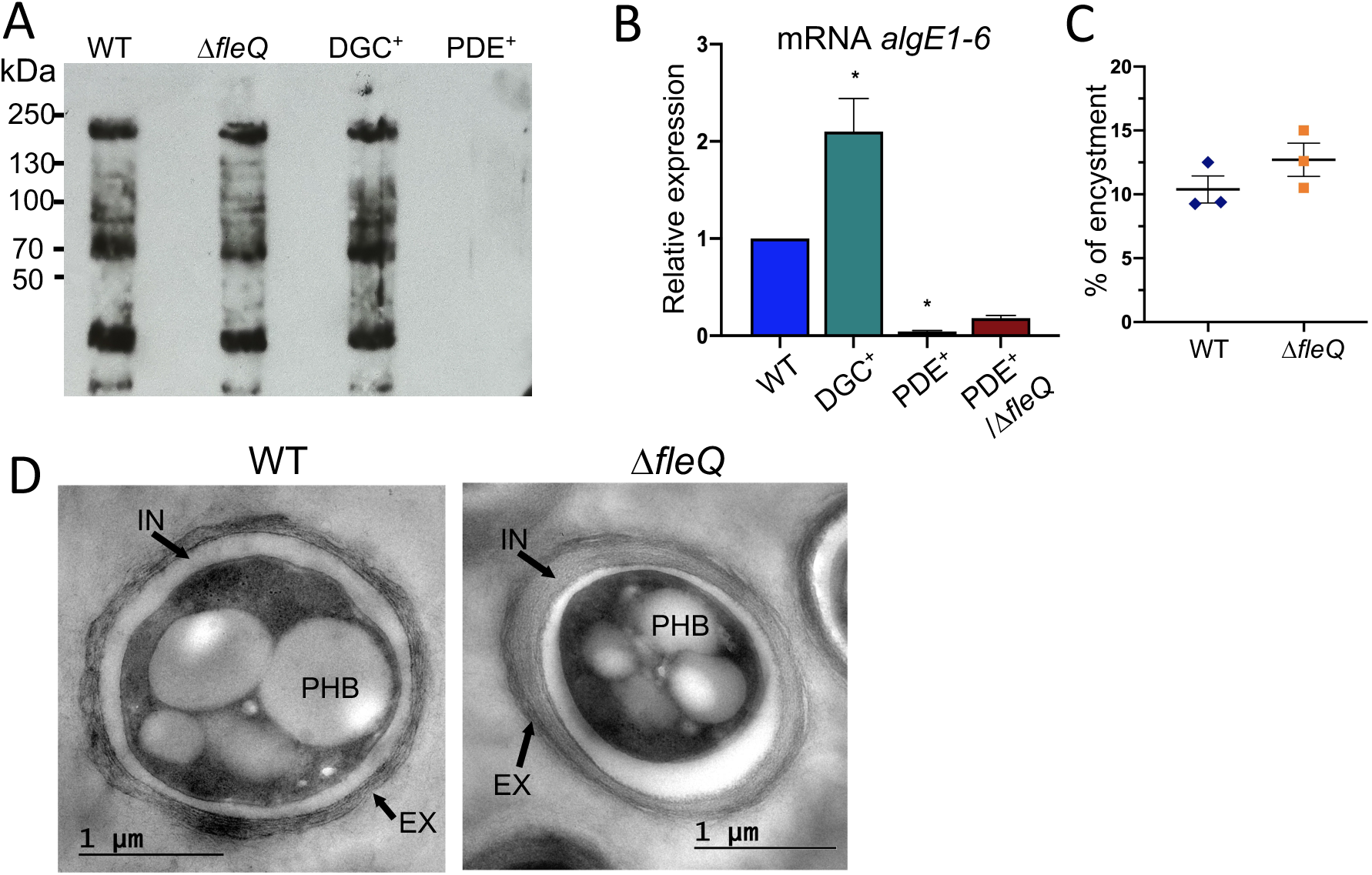
The absence of FleQ does not impair *algE1-6* expression or mature cyst formation. A. Western blot to detect AlgE1-6 epimerases over the surface of wt, Δ*fleQ*, DGC+ or PDE+ differentiated cells, after 5 days on Burk’s-butanol plates. Anti-AlgE4 antibody was used as the primary antibody. B. Determination of the relative abundance of *algE1-6* mRNA by qRT-PCR in cells grown for 48 h in Burk’s-sucrose medium. The bars for standard deviation from three independent experiments are shown. Significant differences were analyzed by *t* test. Statistical significance is shown. *, *P* < 0.01; **, *P* < 0.001. C. Assessment of mature cyst formation based on the percentage of cells resistant to desiccation for 5 days. The bars for standard deviation from three independent experiments (biological replicates) are shown. No significant differences were detected between the two strains. D. Electron micrographs of *A. vinelandii* cysts formed after a 5-day period of differentiation. The cells were induced for encystment on Burk’s-butanol plates. The cyst capsule is clearly observed which is composed of two laminated alginate layers, the exine (EX) and the intine (IN). Poly-B-hydroxybutyrate (PHB) granules are also observed.

### FleQ is not essential for mature cyst formation

Next, we investigated if FleQ could be an intermediary in the regulation of *algE1-6* by c-di-GMP. Western Blot assays revealed that when cultivated under encystment-induced conditions, the Δ*fleQ* mutant showed a pattern of AlgE1-6 detection (Fig 4A) and an accumulation of *algE1-6* transcripts like that of the wt strain (1.11 ± 0.09 for Δ*fleQ* vs 1.0 for wt). These results agreed with a frequency of mature cyst formation by Δ*fleQ* at levels comparable to those shown by wt strain (Fig 4C). The Δ*fleQ* cysts showed a typical morphology, with the exine and intine layers of the alginate protecting coat clearly visible (Fig 4D). These data ruled out a positive essential role of FleQ on the expression of *algE1-6* genes and for the formation of mature cysts.

We also investigated a potential negative role of FleQ on *algE1-6* expression. Low levels of c-di-GMP in strain PDE+ drastically reduced expression of *algE1-6* genes and detection of the corresponding proteins on the surface of the differentiated cell by Western Blot (Fig 4A and B). To reveal a potential repressing effect of FleQ on *algE1-6* expression under reduced c-di-GMP levels, detection of *algE1-6* mRNA was conducted in the double mutant PDE+/Δ*fleQ*. As can be seen in Fig 4B, expression of *algE1-6* genes was not restored in this genetic background, implying that FleQ is not an intermediary in the control of *algE1-6* transcription by the second messenger c-di-GMP.

## Discussion

In contrast to the role of alginate in *P. aeruginosa*, in *A. vinelandii* the alginate conforms the envelope of differentiated cells resistant to desiccation [2]. This fact and the contrasting lifestyles of these two organisms could be the reason of the differing transcriptional regulation of *algD*. In *P. aeruginosa* transcription of the *alg* operon, headed by *algD* is driven by an AlgU-dependent promoter while in *A. vinelandii algD* is transcribed from two well documented sites, from an AlgU and a RpoS promoter [3, 46]. In *P. aeruginosa* several transcriptional factors such as AlgR, AlgB and AmrZ have been reported to directly activate *algD* expression. It has been demonstrated a positive regulatory effect of c-di-GMP on *algD* expression in *P. aeruginosa*. However, the molecular mechanism of such effect remains unknown, as none of the reported transcriptional regulators of *algD* seem to bind c-di-GMP [47].

In this study we demonstrated that *algD* transcription in *A. vinelandii* responds to the intracellular pool of c-di-GMP. Artificially high levels of this second messenger had a positive effect readily detected along the growth curve and it was markedly pronounced during the logarithmic phase of growth (at 12 and 24 h) (Fig 2), while reduced levels of c-di-GMP almost abolished *algD* expression. Given that FleQ is an established c-di-GMP effector in the *Pseudomonadaceae* family, we decided to investigate its influence on *algD* transcription. De-repressed alginate production and *algD* transcription with respect to wt was observed in the Δ*fleQ* mutant, phenotypes that were rescued in the complemented Δ*fleQ*/*fleQ*+ derivative (Fig 1 and 2). Furthermore, a predicted FleQ binding site overlapping the RpoS promoter was identified which was consistent with the ability of FleQ to directly recognize the *algD* regulatory region (Fig 3). Collectively, our data strongly suggest that FleQ represses *algD* expression from the RpoS promoter, contributing to explain the increase in alginate production observed in the Δ*fleQ* mutant. In *P. aeruginosa* at low levels of c-di-GMP FleQ represses transcription of the *pel* operon, whereas at high levels and upon binding to c-di-GMP FleQ activates *pel* transcription [23] At low c-di-GMP levels FleQ binds to two sites (Box 1 and Box2, located 35 bp apart), flanking the −10 and −35 region of the *pelA* promoter and together with FleN promote a distortion of the DNA backbone that prevents the binding of the RNAP [21, 23]. In *A. vinelandii* a second FleQ binding site with a low score (GCCGGCAATGTGAC; p-val 4.73E-4) was also identified 32 bp apart from the first one. Therefore, it is possible to propose a repressing model like that of *P. aeruginosa pelA* for the control of *algD* in *A. vinelandii* under low c-di-GMP levels. This is supported by the fact that a Δ*fleN* mutant of *A. vinelandii* shows an alginate overproducing phenotype that is reflected by the mucoid colony phenotype, when compared to the wt strain (S4 Fig). However, the exact molecular mechanism of the repressing effect of *algD* by FleQ and probably by FleN in *A. vinelandii* needs to be investigated further. It is of interest to note, that reduced levels of c-di-GMP almost abrogated *algD* expression along the growth curve, implying that the AlgU-dependent promoter is also subjected to regulation by the intracellular levels of c-di-GMP as has been reported for *P. aeruginosa* [47].

In the present study we confirmed that transcription of the *algE1-6* genes, encoding alginate C-5 epimerases necessary for the assembly of the cyst capsule, is positively regulated by c-di-GMP. Accumulation of the *algE1-6* mRNA was almost undetectable at low levels of this second messenger relative to the wt strain, which agreed with the lack of detection of the corresponding proteins by Western Blot (Fig 4A and B). In contrast, at high c-di-GMP the *algE1-6* mRNA levels increased by two-fold. However, our data suggest that FleQ is not an intermediary in this regulation since *i*) expression of the *algE1-6* genes was not affected in the Δ*fleQ* mutant and *ii*) expression of *algE1-6* was not restored to wt levels at low c-di-GMP but in the absence of FleQ (Fig 4C). The *algE1-6* genes reside in two regions of the *A. vinelandii* genome (*algE6-4-1-2-3* and *algE5*) and nothing is known about their transcriptional arrangement. It would be interesting to dissect the regulatory mechanism for the control of these genes by c-di-GMP during *A. vinelandii* encystment. Although FleQ was not essential for the formation of mature cysts resistant to desiccation (Fig 4C and D) we cannot rule out a regulatory role of this transcriptional factor during encystment. Work is currently under way to identify additional processes regulated by c-di-GMP and essential for the formation of differentiated cells.

## Acknowledgments

We thank M. C. Gonzaga-Pérez, A. Mejía-Rangel and M. Castañeda-Flores for their technical support; C. A. Gonzalez-Guzmán for his help in protein structure analysis; and G. Zavala for her microscopy technical support. S. Orgaz-Hidalgo was a recipient of a DGAPA-UNAM undergraduate fellowship.

## Supporting information

**S1 Fig. Sequence alignment of FleQ.** FleQ alignment from *A. vinelandii* AEIV (Av) and *P. aeruginosa* PAO1 (P_aer). Motifs for c-di-GMP binding, interaction with sigma 54, or active sites for the AAA+ domain are indicated in red, blue, or green boxes. The HTH domain is highlighted in yellow. Protein alignment was conducted in Clustal Omega (MSA).

**S2 Fig. Alignment of the C-terminal region of FleQ. A**. Structure prediction using the AlphaFold program of the C-terminal region of FleQ from *A. vinelandii* and *P. aeruginosa*, containing the HTH motif. Left panel, superposition of the C-terminus. Right panel, a RMSD value of 0.431 angstroms was obtained considering only the 57 pruned atoms pairs from the HTH domain, using the ChimeraX program. **B.** Sequence alignment of the C-terminal region of FleQ from *A. vinelandii* DJ (AV_DJ), AEIV (AV_AEIV), *Azotobacter Beijerinckia* (A_BEIJ), *Azotobacter croococcum* (A_CRO), *Pseudomonas fluorescens* (P_FL), *Pseudomonas putida* (P_PU), *P. aeruginosa* (P_AER), *Pseudomonas resinovorans* (P_RES), *Pseudomonas otitidis* (P_OT), *Pseudomonas indica* (P_IND). Protein alignment was conducted in Clustal Omega (MSA).

**S3 Fig. *A. vinelandii* FleQ binds to the *P. aeruginosa pelA* promoter**. **A.** DNA sequence of the *pelA* regulatory region. The FleQ binding sites previously reported are in red (Baraquet, et al 2012). The −10 and −35 region of the *pelA* promoter are indicated **B**. EMSA assay to evaluate the binding of *A. vinelandii* FleQ to the *pelA* regulatory region. A DNA fragment was PCR amplified using the primer pair pelA-fw/pelA-Rv. 100 ng of DNA was incubated with increasing concentrations of FleQ. The migration was visualized by staining with ethidium bromide.

**S4 Fig. The D*fleN* mutant shows an alginate-overproducing phenotype.** Growth of the wild type strain (wt) and the D*fleN* mutant on Burk’s-sucrose plates for 48 h. The Δ*fleN* mutant shows an hyper mucoid colony phenotype similar to that of Δ*fleQ*.

**S1 Table.** List of potential FleQ binding sites in the genome of *A. vinelandii* detected by a MEME/MAST analysis using experimentally identified *P. aeruginosa* FleQ binding sites.

