## Supplemental S1-S4 Fig and S1 Table for "The c-di-GMP effector FleQ controls alginate production by repressing transcription of the *algD* gene in *Azotobacter vinelandii*"

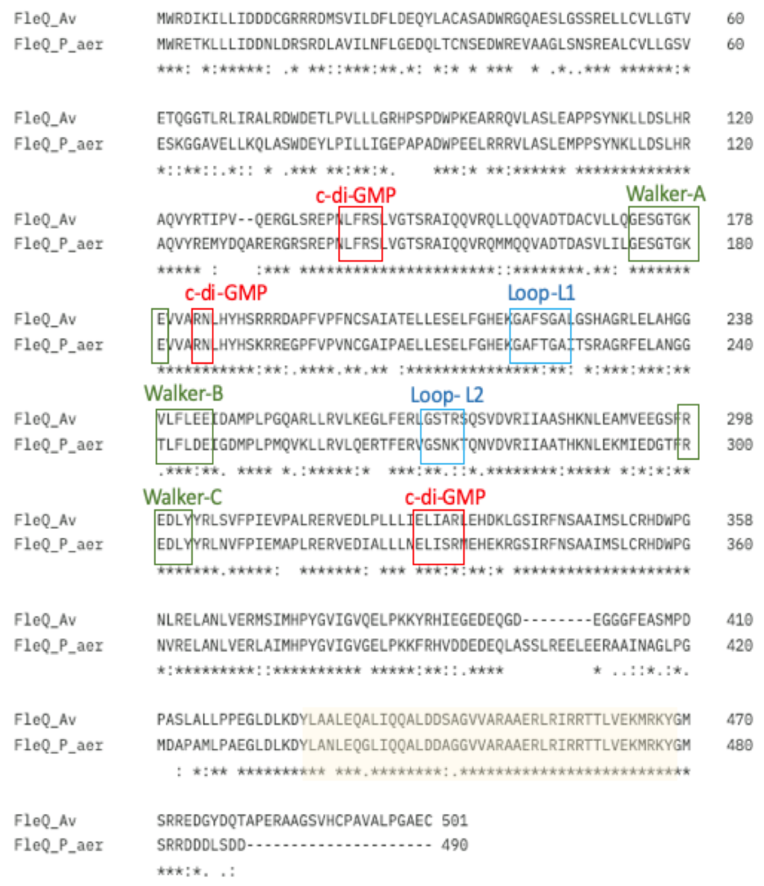

**S1 Fig. Sequence alignment of FleQ. A**. FleQ alignment from *A. vinelandii* AEIV (Av) and *P. aeruginosa* PAO1 (P_aer). Motifs for c-di-GMP binding, interaction with sigma 54, or active sites for the AAA+ domain are indicated in red, blue or green boxes. The HTH domain is highlighted in yellow.

A

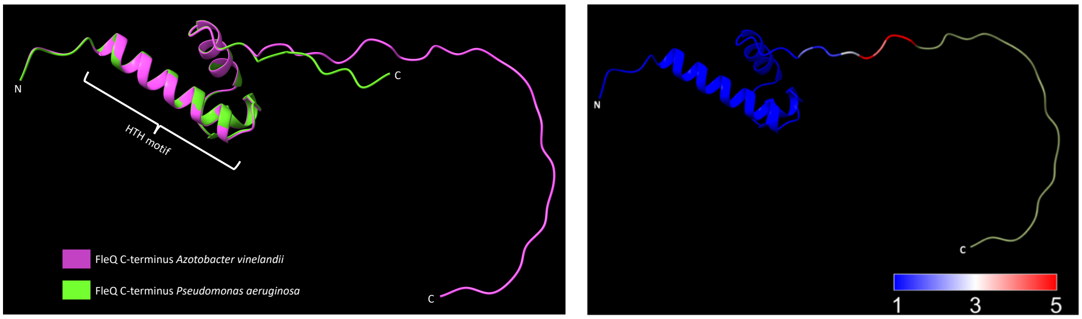

B

```
FLEQ_AV_DJ      PASLALLPPEGLDLKDYLAALEQALIQQALDDSAGVVARAAERLRIRRTTLVEKMRKYGM 470
FLEQ_AV_AEIV    PASLALLPPEGLDLKDYLAALEQALIQQALDDSAGVVARAAERLRIRRTTLVEKMRKYGM 470
FLEQ_A_BEIJ     PSSLALLPPEGLDLKDYLAALEQTLIQQALDDSAGVVARAAERLRIRRTTLVEKMRKYGM 470
FLEQ_A_CRO      PATLALLPPEGLDLKDYLAALEQTLIQQALDDSAGVVARAAERLRIRRTTLVEKMRKYGM 470
FLEQ_P_FL       FSASALLPPEGLDLKDYLGGLEQGLIQQALDDANGIVARAAERLRIRRTTLVEKMRKYGM 480
FLEQ_P_PU       FSNHAMLPPEGLDLKDYLGSLEQGLIQQALDDANGIVARAAERLRIRRTTLVEKMRKYGM 480
FLEQ_P_AER      MDAPAMLPAEGLDLKDYLANLEQGLIQQALDDAGGVVARAAERLRIRRTTLVEKMRKYGM 480
FLEQ_P_RES      IASPALLPPEGLDLKDYLGNLEQGLIQQALDDASGVVARAAERLRIRRTTLVEKMRKYGM 480
FLEQ_P_OT       VSAPAMLPPEGLDLKDYLGNLEQELIQQALDDAAGVVARAAERLRIRRTTLVEKMRKYGM 480
FLEQ_P_IND      MTSPAMLPPEGLDLKDYLGNLEQGLIQQALDDAGGIVARAAERLRIRRTTLVEKMRKYGM 480
                    *:** *********. *** ********: *:************************

FLEQ_AV_DJ      SRREDGYDQTAPERAAGSVHCPAVALPGAEC      501
FLEQ_AV_AEIV    SRREDGYDQTAPERAAGSVHCPAVALPGAEC      501
FLEQ_A_BEIJ     SRREDGGNAVGPGAFVAVDNRSAVAPPGAEC      501
FLEQ_A_CRO      GRREDGDDQPDPESAAAAGHRSAVAPPGAEC      501
FLEQ_P_FL       SRAGGDEQAD---------------------      490
FLEQ_P_PU       SRQGGEDQAED--------------------      491
FLEQ_P_AER      SRRDDDLSDD---------------------      490
FLEQ_P_RES      SRRDEELAEE---------------------      490
FLEQ_P_OT       SRRDEDMAED---------------------      490
FLEQ_P_IND      SRRDDEDQPED--------------------      491
                .*
```

**S2 Fig. Alignment of the C-terminal region of FleQ. A.** Structure prediction using the AlphaFold program of the C-terminal region of FleQ from *A. vinelandii* and *P. aeruginosa*, containing the HTH motif. Left panel, superposition of the C-terminus. Right panel, a RMSD value of 0.431 angstroms was obtained considering only the 57 pruned atoms pairs from the HTH domain. B. Sequence alignment of the C-terminal region of FleQ from *A. vinelandii* DJ (AV_DJ), AEIV (AV_AEIV), *Azotobacter Beijerinckia* (A_BEIJ), *Azotobacter croococcum* (A_CRO), *Pseudomonas fluorescens* (P_FL), *Pseudomonas putida* (P_PU), *P. aeruginosa* (P_AER), *Pseudomonas resinovorans* (P_RES), *Pseudomonas otitidis* (P_OT), *Pseudomonas indica* (P_IND). Protein alignment was conducted in Clustal Omega (MSA).

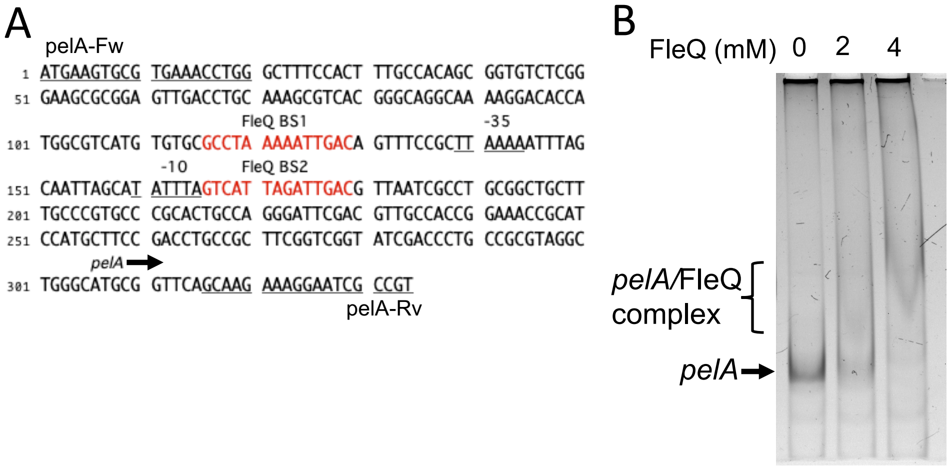

**S3 Fig. *A. vinelandii* FleQ binds to the *P. aeruginosa pelA* promoter. A.** DNA sequence of the *pelA* regulatory region. The FleQ binding sites previously reported are in red (Baraquet, et al 2012). The -10 and -35 region of the *pelA* promoter are indicated **B.** EMSA assay to evaluate the binding of *A. vinelandii* FleQ to the *pelA* regulatory region. A DNA fragment was PCR amplified using the primer pair pelA-fw/pelA-Rv. 100 ng of DNA was incubated with increasing concentrations of FleQ. The migration was visualized by staining with ethidium bromide.

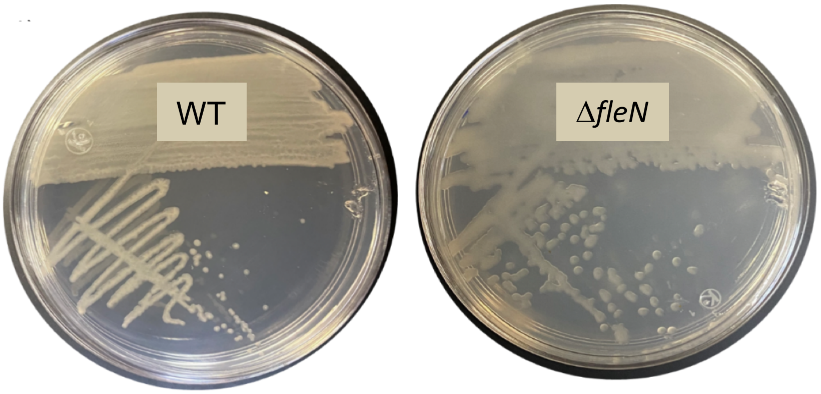

**S4 Fig. The Δ*fleN* mutant shows an alginate-overproducing phenotype.** Growth of the wild type strain (wt) and the Δ*fleN* mutant on Burk's-sucrose plates for 48 h. The Δ*fleN* mutant shows an hyper mucoid colony phenotype similar to that of Δ*fleQ*.

**S1 Table.** List of potential FleQ binding sites in the genome of *A. vinelandii* detected by a MEME/MAST analysis using experimentally identified *P. aeruginosa* FleQ binding sites.

| *locus*_tag | start* | end | Distance to ATG codon | Strand | Score | p-val | q-val | Sequence | Gene | Product |
|---|---|---|---|---|---|---|---|---|---|---|
| Avin_05110 | 490958 | 490971 | 100 | - | 19.4388 | 5.72E-08 | 0.135 | GTCAAAAAACCGAC | | hypothetical protein |
| Avin_05120 | 490958 | 490971 | 349 | - | 19.4388 | 5.72E-08 | 0.123 | GTCAAAAAACCGAC | *ompR* | two-component response regulator OmpR |
| Avin_37690 | 3814816 | 3814829 | -20 | + | 17.2653 | 8.69E-07 | 0.761 | GTCAAAAGATTGGC | | conserved hypothetical protein |
| Avin_50730 | 5141247 | 5141260 | 188 | + | 17.1429 | 9.70E-07 | 0.761 | GTCGGAAAATCGAC | *modA3* | molybdate ABC transporter, periplasmic molybdate-binding protein |
| Avin_52190 | 5334153 | 5334166 | 26 | - | 16.1633 | 2.27E-06 | 1 | GTCGAAAAACTGGC | *atpH* | F1 sector of membrane-bound ATP synthase, delta subunit |
| Avin_12070 | 1160532 | 1160545 | -22 | + | 15.5612 | 3.55E-06 | 1 | GCCAGAAGATCGAC | *pilD* | type IV Pilus Prepilin peptidase, PilD |
| Avin_27160 | 2787068 | 2787081 | -8 | - | 14.7857 | 6.09E-06 | 1 | GTCATTTGATCGAC | | Phytanoyl-CoA dioxygenase |
| Avin_47000 | 4770677 | 4770690 | 224 | - | 14.7245 | 6.37E-06 | 1 | GTCGAAAAACTGTT | *rpoD* | RNA polymerase sigma factor, sigma70; RpoD |
| Avin_08290 | 787011 | 787024 | 27 | + | 14.6429 | 6.70E-06 | 1 | GTCGGAAAACTGGC | | conserved hypothetical protein |
| Avin_08300 | 787011 | 787024 | 227 | + | 14.6429 | 6.70E-06 | 1 | GTCGGAAAACTGGC | *lrgA* | integral membrane protein, LrgA family |
| Avin_00420 | 41396 | 41409 | 355 | - | 14.551 | 7.17E-06 | 1 | GTCGTTATATTGAC | | GGDEF domain protein |
| Avin_17870 | 1774138 | 1774151 | 160 | - | 14.2755 | 8.44E-06 | 1 | GTTATTAGATTGAC | | conserved hypothetical protein |
| Avin_39680 | 4018322 | 4018335 | 190 | - | 14.2653 | 8.47E-06 | 1 | GCCGGAAGATCGAC | | Cytidine/deoxycytidylate deaminase-like protein |
| Avin_27790 | 2862291 | 2862304 | 126 | + | 14.2551 | 8.53E-06 | 1 | GCCGAAAATCCGAC | | UDP-glucose 6-dehydrogenase |
| Avin_27810 | 2862291 | 2862304 | 255 | + | 14.2551 | 8.53E-06 | 1 | GCCGAAAATCCGAC | | hypothetical protein |
| Avin_51880 | 5288801 | 5288814 | 94 | + | 14.1531 | 9.08E-06 | 1 | GCCATAAAATTGAT | | Superfamily I DNA and RNA helicases and helicase subunits-related protein |
| Avin_04180 | 399199 | 399212 | -30 | + | 14.0408 | 9.68E-06 | 1 | GCCAGAAGAGCGAC | | hypothetical protein |
| Avin_04190 | 399199 | 399212 | 141 | + | 14.0408 | 9.68E-06 | 1 | GCCAGAAGAGCGAC | | hypothetical protein |
| Avin_51250 | 5210898 | 5210911 | 186 | - | 13.949 | 1.02E-05 | 1 | GTCGGAAAACTCAC | *algE7* | Secreted bifunctional mannuronan C-5 epimerase/alginate lyase |
| Avin_01710 | 161252 | 161265 | 171 | - | 13.898 | 1.06E-05 | 1 | GTCGCAAATGCGAC | *nifF* | Flavodoxin, nifF |
| Avin_29750 | 3076906 | 3076919 | -17 | - | 13.8367 | 1.10E-05 | 1 | GCCAGAAGTTCGAC | *lpdA* | dihydrolipoamide dehydrogenase |
| Avin_34840 | 3556028 | 3556041 | 105 | + | 13.7653 | 1.14E-05 | 1 | GTCGGAAAACCGGC | | ABC transporter, ATP-binding protein |
| Avin_11960 | 1147671 | 1147684 | 127 | - | 13.6429 | 1.22E-05 | 1 | GCCGAAAAATCGGC | *clpB* | ATP-dependent protease |
| Avin_18580 | 1840518 | 1840531 | 240 | - | 13.5918 | 1.25E-05 | 1 | GTCGGAAAATCCAC | | conserved hypothetical protein |
| Avin_15090 | 1487526 | 1487539 | 383 | + | 13.5714 | 1.27E-05 | 1 | GTCGCAAGATCGGC | *mhpR* | Bacterial regulatory protein, IclR family |
| Avin_18850 | 1872615 | 1872628 | 173 | - | 13.4184 | 1.38E-05 | 1 | GCCAGAAAATCGGC | | conserved hypothetical protein |
| Avin_18860 | 1872615 | 1872628 | -40 | - | 13.4184 | 1.38E-05 | 1 | GCCAGAAAATCGGC | | NAD-dependent glutamate dehydrogenase |
| Avin_39180 | 3970876 | 3970889 | 205 | + | 13.3878 | 1.40E-05 | 1 | GTCGAAAATTCCAC | *cspA* | Cold-shock-like protein |
| Avin_39190 | 3970876 | 3970889 | -22 | + | 13.3878 | 1.40E-05 | 1 | GTCGAAAATTCCAC | | hypothetical protein |
| Avin_60180 | 1995846 | 1995859 | 112 | - | 13.051 | 1.68E-05 | 1 | GTCGCAAGACCGGC | *tRNA* | tRNA-Asn |
| Avin_11330 | 1080697 | 1080710 | -27 | - | 13.0408 | 1.69E-05 | 1 | GTCGATTGTCCGAC | | hypothetical protein |
| Avin_11340 | 1080697 | 1080710 | 51 | - | 13.0408 | 1.69E-05 | 1 | GTCGATTGTCCGAC | | Bacterial outer membrane porin |
| Avin_27550 | 2833221 | 2833234 | -40 | + | 12.898 | 1.82E-05 | 1 | GCCAGAAAACCGGC | | Peptidase, S49 family |
| Avin_27570 | 2833221 | 2833234 | 216 | + | 12.898 | 1.82E-05 | 1 | GCCAGAAAACCGGC | | fructose-2,6-bisphosphatase |
| Avin_16040 | 1585672 | 1585685 | 357 | - | 12.8265 | 1.89E-05 | 1 | GTCACTATTTTGTC | | hypothetical protein |
| Avin_16050 | 1585672 | 1585685 | 157 | - | 12.8265 | 1.89E-05 | 1 | GTCACTATTTTGTC | | conserved hypothetical protein |
| Avin_16060 | 1585672 | 1585685 | 360 | - | 12.8265 | 1.89E-05 | 1 | GTCACTATTTTGTC | | hypothetical protein |
| Avin_19220 | 1917177 | 1917190 | 129 | + | 12.6735 | 2.03E-05 | 1 | GTCGAAAGTTCCAC | *rnfE* | Electron transport complex, subunit E |
| Avin_35450 | 3618054 | 3618067 | 330 | - | 12.6224 | 2.09E-05 | 1 | GTCGATAAAGCGTT | | hypothetical protein |
| Avin_23460 | 2348107 | 2348120 | 81 | - | 12.5408 | 2.17E-05 | 1 | GCCGAAAGACCGAT | *nasS* | nitrate/nitrite transport system substrate-binding protein ; NasS |
| Avin_35250 | 3602312 | 3602325 | 199 | - | 12.4898 | 2.23E-05 | 1 | GTCGGAAGACTGTT | | hypothetical protein |
| Avin_35260 | 3602312 | 3602325 | 15 | - | 12.4898 | 2.23E-05 | 1 | GTCGGAAGACTGTT | | hypothetical protein |
| Avin_35270 | 3602312 | 3602325 | 238 | - | 12.4898 | 2.23E-05 | 1 | GTCGGAAGACTGTT | | conserved hypothetical protein |
| Avin_44590 | 4509487 | 4509500 | 286 | + | 12.4796 | 2.24E-05 | 1 | GCCGGAAAACTGGC | | conserved hypothetical protein |
| Avin_39490 | 4000343 | 4000356 | 231 | - | 12.4286 | 2.29E-05 | 1 | GTCAGATGATCGGC | *hom* | homoserine dehydrogenase |
| Avin_16060 | 1585949 | 1585962 | 83 | - | 12.4082 | 2.32E-05 | 1 | GCCGAAAGACCGGC | | hypothetical protein |
| Avin_30500 | 3160798 | 3160811 | 50 | - | 12.398 | 2.33E-05 | 1 | GTCAGTAGTCCGGC | *ssuD* | alkanesulfonate monooxygenase |
| Avin_51880 | 5288681 | 5288694 | -26 | + | 12.3367 | 2.41E-05 | 1 | GTCTAAATAGCGAC | | Superfamily I DNA and RNA helicases and helicase subunits-related protein |
| Avin_30380 | 3150289 | 3150302 | 66 | - | 12.2653 | 2.49E-05 | 1 | GCCACAAGTTCGTC | *ccmH* | Cytochrome C biogenesis protein |
| Avin_49000 | 4961202 | 4961215 | 148 | - | 12.2551 | 2.50E-05 | 1 | GCCGGTAAACTGGC | *anfH* | nitrogenase iron protein |
| Avin_04470 | 424288 | 424301 | 129 | - | 12.2041 | 2.56E-05 | 1 | GTCAGTTGATCGGC | *cooC* | Carbon monoxide dehydrogenae assesory protein, CooC |
| Avin_25460 | 2554247 | 2554260 | 158 | - | 12.2041 | 2.56E-05 | 1 | GTCGGAAAAGTGTT | | hypothetical protein |
| Avin_48870 | 4947131 | 4947144 | 146 | + | 12.1224 | 2.66E-05 | 1 | GCCGGAAAATCGGC | *spuA* | glutamine amidotransferase |
| Avin_48880 | 4947131 | 4947144 | 117 | + | 12.1224 | 2.66E-05 | 1 | GCCGGAAAATCGGC | | glutamine synthetase |

| | | | | | | | | | | |
|---|---|---|---|---|---|---|---|---|---|---|
| Avin 25210 | 2527539 | 2527552 | 92 | - | 12.051 | 2.75E-05 | 1 | GTCGCAAGAGCGGC | | acyl-CoA dehydrogenase |
| Avin 00370 | 38375 | 38388 | 246 | + | 12.0408 | 2.76E-05 | 1 | GTCGGAAATCCGGC | | transmembrane protein with C-terminal GGDEF motif |
| Avin 08030 | 759646 | 759659 | 13 | - | 12.0306 | 2.78E-05 | 1 | GCCAGAAAAGCGAT | *mpl* | UDP-N-acetylmuramate:L-alanyl-gamma-D-glutamyl-meso-diaminopimelate lig |
| Avin 37860 | 3829822 | 3829835 | 298 | + | 12.0102 | 2.80E-05 | 1 | GCCAGAAGATCCAC | | conserved hypothetical protein |
| Avin 01770 | 166897 | 166910 | 142 | - | 11.9286 | 2.91E-05 | 1 | GTCTGTTGATCGAC | | conserved hypothetical protein |
| Avin 15680 | 1537867 | 1537880 | 146 | + | 11.9082 | 2.94E-05 | 1 | GCCTCAAGACCGAC | *edd-1* | 6-phosphogluconate dehydratase |
| Avin 38290 | 3874472 | 3874485 | 210 | + | 11.898 | 2.96E-05 | 1 | GCCACAAAAGCGGC | | hypothetical protein |
| Avin 38310 | 3874472 | 3874485 | 110 | + | 11.898 | 2.96E-05 | 1 | GCCACAAAAGCGGC | *mvaT* | Transcriptional regulator MvaT |
| Avin 35460 | 3619546 | 3619559 | 371 | - | 11.8878 | 2.97E-05 | 1 | GCCGGAAAAGCGTC | | hypothetical protein |
| Avin 46200 | 4695867 | 4695880 | 146 | - | 11.8673 | 3.00E-05 | 1 | GTCACATGATTGTT | | conserved hypothetical protein |
| Avin 03960 | 377973 | 377986 | 275 | + | 11.8469 | 3.02E-05 | 1 | GCCGGAAGTTTGTC | | conserved hypothetical protein |
| Avin 60150 | 1678775 | 1678788 | -27 | + | 11.8265 | 3.05E-05 | 1 | GTCGGTAGAGCGGC | *tRNA* | tRNA-Thr |
| Avin 24310 | 2424232 | 2424245 | 25 | - | 11.8265 | 3.05E-05 | 1 | GCCGCATGATCGAC | *flhC* | Flagellar transcriptional activator |
| Avin 27020 | 2771872 | 2771885 | 189 | + | 11.8265 | 3.05E-05 | 1 | GCCGGATGATCGAC | | hypothetical protein |
| Avin 38420 | 3887445 | 3887458 | 29 | - | 11.8265 | 3.05E-05 | 1 | GCCGGTAGAGTGTC | | GGDEF Response Regulator |
| Avin 38440 | 3887445 | 3887458 | 150 | - | 11.8265 | 3.05E-05 | 1 | GCCGGTAGAGTGTC | *fumB* | fumarate hydratase, class I |
| Avin 20420 | 2034191 | 2034204 | 55 | - | 11.8061 | 3.08E-05 | 1 | GCCAGTAAAGCGAT | *thrS* | threonyl-tRNA synthetase |
| Avin 05070 | 481496 | 481509 | 354 | - | 11.7857 | 3.10E-05 | 1 | GCCTCAAGAGTGAC | *bcsD* | Cellulose synthase subunit D |
| Avin 00180 | 20220 | 20233 | 237 | - | 11.602 | 3.38E-05 | 1 | GCCGGTTGATCGAC | *lysM* | peptidoglycan-binding LysM protein |
| Avin 43940 | 4442339 | 4442352 | 243 | - | 11.5918 | 3.39E-05 | 1 | GTCGATAATTCCTC | | TonB-dependent receptor family |
| Avin 29140 | 3006175 | 3006188 | 152 | - | 11.551 | 3.44E-05 | 1 | GCCAGATGATCGTC | | conserved hypothetical protein |
| Avin 28760 | 2961921 | 2961934 | 176 | + | 11.4898 | 3.55E-05 | 1 | GCCGAAAGATCGTT | *nfuA* | NfuA protein |
| Avin 21990 | 2198432 | 2198445 | 175 | - | 11.4388 | 3.63E-05 | 1 | GTCTAAATTTTGTC | | ABC transporter, inner membrane permease component |
| Avin 22000 | 2198432 | 2198445 | 19 | - | 11.4388 | 3.63E-05 | 1 | GTCTAAATTTTGTC | | transcriptional regulator, Crp/Fnr family |
| Avin 18540 | 1833471 | 1833484 | 167 | - | 11.4082 | 3.68E-05 | 1 | GCCGGAAGATCGGC | | outer membrane efflux protein |
| Avin 20010 | 1989057 | 1989070 | 195 | + | 11.3673 | 3.74E-05 | 1 | GTCTCAAAACTGTT | *ccoN* | cytochrome c oxidase, cbb3-type, subunit I |
| Avin 20020 | 1989057 | 1989070 | 217 | + | 11.3673 | 3.74E-05 | 1 | GTCTCAAAACTGTT | | conserved hypothetical protein |
| Avin 05330 | 513845 | 513858 | -21 | + | 11.2653 | 3.90E-05 | 1 | GCCAGAAGATCGTT | | Glycosyl transferase, family 2 |
| Avin 40410 | 4092022 | 4092035 | -6 | + | 11.2653 | 3.90E-05 | 1 | GTCGCATGATCGAT | *iscR* | Iron-sulphur cluster assembly transcription factor IscR |
| Avin 45300 | 4596506 | 4596519 | 85 | + | 11.1939 | 4.03E-05 | 1 | GCCTATAATGCGAC | *pilM* | Type IV pilus assembly protein |
| Avin 45310 | 4596506 | 4596519 | 66 | + | 11.1939 | 4.03E-05 | 1 | GCCTATAATGCGAC | *ponA* | Penicillin binding protein 1A |
| Avin 23100 | 2306521 | 2306534 | -49 | + | 11.1837 | 4.05E-05 | 1 | GCCAGAAGAGCGGC | *oprI* | outer membrane lipoprotein OprI |
| Avin 34230 | 3497819 | 3497832 | 316 | - | 11.1327 | 4.12E-05 | 1 | GCCGGATGATTGTC | | conserved hypothetical protein |
| Avin 14050 | 1373815 | 1373828 | 95 | + | 11.1224 | 4.14E-05 | 1 | GTCTCTAGTGCGAC | *bfr* | bacterioferritin |
| Avin 26730 | 2743375 | 2743388 | 208 | + | 11.0816 | 4.22E-05 | 1 | GTCACATATCTCAC | | conserved hypothetical protein |
| Avin 48350 | 4900880 | 4900893 | 62 | - | 11.0714 | 4.24E-05 | 1 | GTCGATAATCCCTC | | ammonia monooxygenase |
| Avin 10970 | 1050741 | 1050754 | 101 | + | 11.0408 | 4.30E-05 | 1 | GTCAATAAAATGTC | *algD* | GDP-mannose 6-dehydrogenase |
| Avin 12680 | 1232969 | 1232982 | 14 | - | 11.0408 | 4.30E-05 | 1 | GTCTGAAGATCCAC | *maf* | septum formation protein |
| Avin 40650 | 4108261 | 4108274 | 29 | - | 11.0204 | 4.34E-05 | 1 | GTTGAAAATCCGTC | | hypothetical protein |
| Avin 40660 | 4108261 | 4108274 | 384 | - | 11.0204 | 4.34E-05 | 1 | GTTGAAAATCCGTC | | hypothetical protein |
| Avin 06280 | 608611 | 608624 | 139 | + | 11 | 4.38E-05 | 1 | GTCGCTAATGTCAC | *rplB* | 50S ribosomal protein L2 |
| Avin 13260 | 1289828 | 1289841 | 180 | - | 10.9796 | 4.40E-05 | 1 | GCCAGATAATCGGC | *murC* | UDP-N-acetylmuramate--alanine ligase |
| Avin 35200 | 3594005 | 3594018 | 232 | + | 10.9694 | 4.42E-05 | 1 | GTCAATAGATTGAA | | conserved hypothetical protein |
| Avin 55010 | 179363 | 179376 | 114 | + | 10.8878 | 4.60E-05 | 1 | GTCGATTGTTTCAC | *rRNA* | ribosomal RNA(operon 1/6) |
| Avin 55040 | 1379899 | 1379912 | 114 | + | 10.8878 | 4.60E-05 | 1 | GTCGATTGTTTCAC | *rRNA* | ribosomal RNA(operon 2/6) |
| Avin 55070 | 1911170 | 1911183 | 114 | + | 10.8878 | 4.60E-05 | 1 | GTCGATTGTTTCAC | *rRNA* | ribosomal RNA(operon 3/6) |
| Avin 55100 | 2840775 | 2840788 | 114 | - | 10.8878 | 4.60E-05 | 1 | GTCGATTGTTTCAC | *rRNA* | ribosomal RNA(operon 4/6) |
| Avin 55130 | 4279011 | 4279024 | 114 | - | 10.8878 | 4.60E-05 | 1 | GTCGATTGTTTCAC | *rRNA* | ribosomal RNA(operon 5/6) |
| Avin 55160 | 4672946 | 4672959 | 114 | - | 10.8878 | 4.60E-05 | 1 | GTCGATTGTTTCAC | *rRNA* | ribosomal RNA(operon 6/6) |
| Avin 05450 | 523825 | 523838 | 285 | - | 10.8163 | 4.73E-05 | 1 | GTTGCAAAACTGGC | *pckA* | phosphoenolpyruvate carboxykinase |
| Avin 29700 | 3069969 | 3069982 | 94 | + | 10.7959 | 4.78E-05 | 1 | GCTGGAAGACTGAC | | Xanthine/uracil permease family |
| Avin 50730 | 5141247 | 5141260 | 188 | - | 10.7857 | 4.79E-05 | 1 | GTCGATTTTCCGAC | *modA3* | molybdate ABC transporter, periplasmic molybdate-binding protein |
| Avin 50210 | 5090337 | 5090350 | 126 | - | 10.7041 | 4.95E-05 | 1 | GTCGATTGTTCGGC | | LamB type porin |
| Avin 51410 | 5226642 | 5226655 | 20 | + | 10.7041 | 4.95E-05 | 1 | GTCGATTGTTCGGC | | Glycoside hydrolase, clan GH-D |
| Avin 05110 | 490958 | 490971 | 100 | + | 10.6633 | 5.04E-05 | 1 | GTCGGTTTTTTGAC | | hypothetical protein |
| Avin 05120 | 490958 | 490971 | 349 | + | 10.6633 | 5.04E-05 | 1 | GTCGGTTTTTTGAC | *ompR* | two-component response regulator OmpR |
| Avin 21240 | 2122413 | 2122426 | 54 | - | 10.6429 | 5.08E-05 | 1 | GTCGAAAGTTCGGT | | hypothetical protein |
| Avin 40100 | 4066070 | 4066083 | 173 | - | 10.602 | 5.17E-05 | 1 | GTCTGAAAATTGGT | | hypothetical protein |
| Avin 40120 | 4066070 | 4066083 | 111 | - | 10.602 | 5.17E-05 | 1 | GTCTGAAAATTGGT | | 2-isopropylmalate synthase |

| | | | | | | | | | | |
|---|---|---|---|---|---|---|---|---|---|---|
| Avin 17200 | 1704303 | 1704316 | 272 | + | 10.4898 | 5.40E-05 | 1 | GTCGCATGAGTGGC | | CRISPR-associated protein, CT1975 |
| Avin 31670 | 3277399 | 3277412 | 267 | - | 10.449 | 5.49E-05 | 1 | GCCAATAATTTCAT | *oprE* | outer membrane porin OprE |
| Avin 31680 | 3277399 | 3277412 | 121 | - | 10.449 | 5.49E-05 | 1 | GCCAATAATTTCAT | | ABC transporter, aliphatic sulfonate substrate-binding protein |
| Avin 02830 | 268816 | 268829 | 345 | - | 10.4388 | 5.51E-05 | 1 | GCCAGAAGATCCTC | *gmk* | guanylate kinase |
| Avin 46860 | 4755432 | 4755445 | 230 | + | 10.4388 | 5.51E-05 | 1 | GCCAGAAGATCCTC | *glpE* | Thiosulfate sulfurtransferase |
| Avin 25090 | 2511879 | 2511892 | 280 | - | 10.4082 | 5.57E-05 | 1 | GTCGAAAGTGCGTT | | Major facilitator superfamily transporter |
| Avin 38800 | 3923961 | 3923974 | 337 | + | 10.3776 | 5.66E-05 | 1 | GTCGGAAGACTCGC | *kdsA* | 3-deoxy-8-phosphooctulonate synthase |
| Avin 28700 | 2957096 | 2957109 | 298 | - | 10.3469 | 5.73E-05 | 1 | GTCGGAAAACCCAT | | hypothetical protein |
| Avin 31060 | 3209593 | 3209606 | 201 | - | 10.3061 | 5.83E-05 | 1 | GCCGGATGAGCGAC | | hypothetical protein |
| Avin 35760 | 3648691 | 3648704 | 384 | + | 10.3061 | 5.83E-05 | 1 | GCCACAAATGCGAT | | Bacterial regulatory protein, LysR family |
| Avin 11650 | 1121008 | 1121021 | 364 | - | 10.2653 | 5.91E-05 | 1 | GCCAGATATTTGAT | *pepA* | Leucyl aminopeptidase |
| Avin 40980 | 4135416 | 4135429 | 203 | + | 10.2551 | 5.93E-05 | 1 | GCCAAATATCCGGC | *cobW* | Cobalamin synthesis protein |
| Avin 40990 | 4135416 | 4135429 | -20 | + | 10.2551 | 5.93E-05 | 1 | GCCAAATATCCGGC | | hypothetical protein |
| Avin 36530 | 3715697 | 3715710 | 51 | + | 10.2449 | 5.96E-05 | 1 | GCCAAAATTTCGGC | | hypothetical protein |
| Avin 36540 | 3715697 | 3715710 | 194 | + | 10.2449 | 5.96E-05 | 1 | GCCAAAATTTCGGC | | Fic (filamentation induced by cAMP) protein |
| Avin 16280 | 1612680 | 1612693 | 34 | - | 10.1939 | 6.09E-05 | 1 | GTTGATAGTTTGGC | | Prevent-host-death protein |
| Avin 16290 | 1612680 | 1612693 | 240 | - | 10.1939 | 6.09E-05 | 1 | GTTGATAGTTTGGC | | addiction module toxin, Txe/YoeB family |
| Avin 08610 | 818228 | 818241 | 233 | + | 10.1633 | 6.17E-05 | 1 | GCTACAAGATCGTC | *benC* | Benzoate 1,2-Dioxygenase Reductase |
| Avin 44610 | 4514066 | 4514079 | 87 | + | 10.0816 | 6.38E-05 | 1 | GGCAAAAGACCGAC | | 3-deoxy-D-manno-octulosonic-acid transferase |
| Avin 44620 | 4514066 | 4514079 | 144 | + | 10.0816 | 6.38E-05 | 1 | GGCAAAAGACCGAC | | FAD dependent oxidoreductase |
| Avin 34150 | 3489170 | 3489183 | -8 | - | 10.0612 | 6.43E-05 | 1 | GTCACATGTGTGGC | *purF* | Amidophosphoribosyl transferase |
| Avin 01760 | 165769 | 165782 | 175 | - | 10.0408 | 6.47E-05 | 1 | GCCAGTTGATCGGC | | Thioesterase superfamily protein |
| Avin 44140 | 4468473 | 4468486 | 294 | - | 9.9898 | 6.60E-05 | 1 | GCCTGATGATCGAC | | PupR/FecR-like anti-sigma factor protein |
| Avin 31960 | 3297143 | 3297156 | -17 | + | 9.96939 | 6.66E-05 | 1 | GTCGAAAAACTTAC | | transcriptional regulator protein |
| Avin 45290 | 4595316 | 4595329 | -22 | + | 9.95918 | 6.69E-05 | 1 | GGCAGAAGATTGAC | *pilN* | Fimbrial assembly protein |
| Avin 32050 | 3304564 | 3304577 | 214 | - | 9.94898 | 6.72E-05 | 1 | GTCGCTTGTCCGTC | | Conserved hypothetical protein |
| Avin 32060 | 3304564 | 3304577 | 84 | - | 9.94898 | 6.72E-05 | 1 | GTCGCTTGTCCGTC | | hypothetical protein |
| Avin 49000 | 4961383 | 4961396 | 329 | - | 9.94898 | 6.72E-05 | 1 | GTCGGTAAATTCTT | *anfH* | nitrogenase iron protein |
| Avin 43060 | 4345553 | 4345566 | 216 | - | 9.90816 | 6.83E-05 | 1 | GTTGATAAATTGGT | | GntR family transcriptional regulator |
| Avin 43070 | 4345553 | 4345566 | 126 | - | 9.90816 | 6.83E-05 | 1 | GTTGATAAATTGGT | *lctP* | L-lactate permease |
| Avin 16660 | 1650519 | 1650532 | 42 | + | 9.88776 | 6.89E-05 | 1 | GTCATTAAATTGAG | | ISRSO17-transposase protein |
| Avin 22400 | 2241291 | 2241304 | 47 | + | 9.88776 | 6.89E-05 | 1 | GCCGGAAGAGCGGC | *dszA* | xenobiotic compound monooxygenase, DszA family, A subunit protein |
| Avin 22410 | 2241291 | 2241304 | 19 | + | 9.88776 | 6.89E-05 | 1 | GCCGGAAGAGCGGC | | hypothetical protein |
| Avin 22420 | 2241291 | 2241304 | 354 | + | 9.88776 | 6.89E-05 | 1 | GCCGGAAGAGCGGC | | ABC nitrate/sulfonate/bicarbonate family transporter protein |
| Avin 32020 | 3300906 | 3300919 | 54 | + | 9.88776 | 6.89E-05 | 1 | GTCATTAAATTGAG | | conserved hypothetical protein |
| Avin 32030 | 3300906 | 3300919 | 42 | + | 9.88776 | 6.89E-05 | 1 | GTCATTAAATTGAG | | ISRSO17-transposase protein |
| Avin 50370 | 5107699 | 5107712 | 112 | - | 9.88776 | 6.89E-05 | 1 | GTCGGATATGCGTC | | Aspartate racemase |
| Avin 13440 | 1310352 | 1310365 | 169 | - | 9.83673 | 7.02E-05 | 1 | GCCGAATATCTGGC | *pdxH* | pyridoxamine 5-phosphate oxidase |
| Avin 20370 | 2027678 | 2027691 | 360 | + | 9.82653 | 7.04E-05 | 1 | GCCAGATGTTCGTC | | hypothetical protein |
| Avin 36530 | 3715697 | 3715710 | 51 | - | 9.82653 | 7.04E-05 | 1 | GCCGAAATTTTGGC | | hypothetical protein |
| Avin 36540 | 3715697 | 3715710 | 194 | - | 9.82653 | 7.04E-05 | 1 | GCCGAAATTTTGGC | | Fic (filamentation induced by cAMP) protein |
| Avin 21480 | 2148135 | 2148148 | -28 | + | 9.81633 | 7.06E-05 | 1 | GTCGGTAAAGCGGT | | conserved hypothetical protein |
| Avin 06770 | 642530 | 642543 | 263 | + | 9.79592 | 7.13E-05 | 1 | GTCAAAAGACCGAG | *thiL* | thiamine-monophosphate kinase |
| Avin 25900 | 2647458 | 2647471 | 309 | - | 9.77551 | 7.18E-05 | 1 | GCCAGTAAATCCAT | | conserved hypothetical protein |
| Avin 12400 | 1205516 | 1205529 | 22 | - | 9.76531 | 7.20E-05 | 1 | GCCGGAATTCCGAC | | PepSY-associated TM helix |
| Avin 01370 | 136470 | 136483 | 165 | - | 9.7449 | 7.27E-05 | 1 | GTGACAAAACTGAC | | conserved hypothetical protein |
| Avin 01380 | 136470 | 136483 | 276 | - | 9.7449 | 7.27E-05 | 1 | GTGACAAAACTGAC | *nifH* | Nitrogenase iron protein |
| Avin 50680 | 5137060 | 5137073 | 126 | - | 9.7449 | 7.27E-05 | 1 | GTCGTAAAAATGAC | *modE* | Mo regulation, Mo processing homeostasis |
| Avin 50690 | 5137060 | 5137073 | 67 | - | 9.7449 | 7.27E-05 | 1 | GTCGTAAAAATGAC | *modG* | Mo processing, homeostasis |
| Avin 22180 | 2217765 | 2217778 | 71 | - | 9.72449 | 7.34E-05 | 1 | GTCGATAGACTGCC | | hypothetical protein |
| Avin 22190 | 2217765 | 2217778 | 52 | - | 9.72449 | 7.34E-05 | 1 | GTCGATAGACTGCC | *pykA-2* | Pyruvate kinase |
| Avin 34320 | 3506487 | 3506500 | 272 | - | 9.72449 | 7.34E-05 | 1 | GTCACGAGATCGAC | *dusC* | tRNA-dihydrouridine synthase C |
| Avin 48970 | 4958268 | 4958281 | 59 | + | 9.72449 | 7.34E-05 | 1 | GTCAGATAATCCAT | *anfK* | Fe-only nitrogenase, beta subunit |
| Avin 16360 | 1618707 | 1618720 | 385 | - | 9.68367 | 7.44E-05 | 1 | GCTAAATAATTGAT | | major facilitator family transporter |
| Avin 13260 | 1289828 | 1289841 | 180 | + | 9.61224 | 7.66E-05 | 1 | GCCGATTATCTGGC | *murC* | UDP-N-acetylmuramate--alanine ligase |
| Avin 06960 | 661661 | 661674 | -28 | - | 9.59184 | 7.71E-05 | 1 | GTCAAGAAACTGAT | *accB* | acetyl-CoA carboxylase, biotin carboxyl carrier protein |
| Avin 02470 | 237126 | 237139 | -20 | - | 9.58163 | 7.73E-05 | 1 | GTCGCAAGTTCCTC | *aglA-2* | Alpha-glucosidase |
| Avin 41820 | 4218471 | 4218484 | -13 | + | 9.55102 | 7.83E-05 | 1 | GTCACAATACTCAT | | Lysine 2,3-aminomutase |
| Avin 41840 | 4218471 | 4218484 | 129 | + | 9.55102 | 7.83E-05 | 1 | GTCACAATACTCAT | | Bacterial regulatory protein, LysR family |

| | | | | | | | | | | |
|---|---|---|---|---|---|---|---|---|---|---|
| Avin 03160 | 295073 | 295086 | -1 | - | 9.5102 | 7.96E-05 | 1 | GCCGAAAGATCCAT | | TonB protein |
| Avin 43160 | 4359155 | 4359168 | 23 | + | 9.5102 | 7.96E-05 | 1 | GCCAGTATTTTGAT | | Resolvase-like protein |
| Avin 13800 | 1341060 | 1341073 | -25 | + | 9.47959 | 8.06E-05 | 1 | GCCGAATATTCGGC | | hypothetical protein |
| Avin 13800 | 1341060 | 1341073 | -25 | - | 9.47959 | 8.06E-05 | 1 | GCCGAATATTCGGC | | hypothetical protein |
| Avin 25390 | 2546821 | 2546834 | 21 | + | 9.46939 | 8.09E-05 | 1 | GCCTGATGACCGAC | *fpvI* | RNA polymerase sigma factor, FecI family |
| Avin 25400 | 2546821 | 2546834 | 136 | + | 9.46939 | 8.09E-05 | 1 | GCCTGATGACCGAC | *fpvR* | Anti-sigma factor, FecR family |
| Avin 26970 | 2764833 | 2764846 | 260 | - | 9.46939 | 8.09E-05 | 1 | GTCTTTATTGTGAC | | conserved hypothetical protein-transmembrane prediction |
| Avin 13600 | 1326076 | 1326089 | 245 | - | 9.45918 | 8.12E-05 | 1 | GCCACAAGTGCGGC | | transposase |
| Avin 13610 | 1326076 | 1326089 | 64 | - | 9.45918 | 8.12E-05 | 1 | GCCACAAGTGCGGC | *ung* | uracil-DNA glycosylase |
| Avin 45270 | 4594080 | 4594093 | 15 | + | 9.42857 | 8.23E-05 | 1 | GTCGTTATAGCGAT | *pilP* | Pilus assembly protein |
| Avin 48670 | 4929553 | 4929566 | 130 | + | 9.40816 | 8.27E-05 | 1 | GTCACAAAGCCGAC | | conserved hypothetical protein |
| Avin 48680 | 4929553 | 4929566 | 117 | + | 9.40816 | 8.27E-05 | 1 | GTCACAAAGCCGAC | | mcbC-like oxidoreductase |
| Avin 16480 | 1633994 | 1634007 | 135 | - | 9.38776 | 8.36E-05 | 1 | GTCAAAAAAGCAAC | | ROK-related protein |
| Avin 16280 | 1612585 | 1612598 | 129 | - | 9.35714 | 8.45E-05 | 1 | GCCGCTTGTCCGAC | | Prevent-host-death protein |
| Avin 16290 | 1612585 | 1612598 | 335 | - | 9.35714 | 8.45E-05 | 1 | GCCGCTTGTCCGAC | | addiction module toxin, Txe/YoeB family |
| Avin 23130 | 2308651 | 2308664 | 213 | + | 9.35714 | 8.45E-05 | 1 | GCTGATAAAGCGTC | *aroG* | Phospho-2-dehydro-3-deoxyheptonate aldolase, subtype 1 |
| Avin 23140 | 2308651 | 2308664 | 86 | + | 9.35714 | 8.45E-05 | 1 | GCTGATAAAGCGTC | *cysB* | LysR family transcriptional regulator protein |
| Avin 33500 | 3425509 | 3425522 | -23 | + | 9.35714 | 8.45E-05 | 1 | GTCGGAATTGCGTC | | esterase, poly(3-hydroxybutyrate) depolymerase |
| Avin 65010 | 397467 | 397480 | -45 | - | 9.30612 | 8.61E-05 | 1 | GTCGAAGGATCGAC | *ncRNA* | PrrB RsmZ sRNA family; predicted by Infernal |
| Avin 65020 | 400438 | 400451 | -45 | - | 9.30612 | 8.61E-05 | 1 | GTCGAAGGATCGAC | *ncRNA* | PrrB RsmZ sRNA family; predicted by Infernal |
| Avin 08930 | 844018 | 844031 | 298 | - | 9.30612 | 8.61E-05 | 1 | GTCGAAGGATCGAC | | conserved hypothetical protein |
| Avin 08950 | 844018 | 844031 | 188 | - | 9.30612 | 8.61E-05 | 1 | GTCGAAGGATCGAC | | Staphylococcus nuclease (SNase-like) |
| Avin 15420 | 1515400 | 1515413 | -19 | - | 9.28571 | 8.68E-05 | 1 | GTCAGTAAATCGAA | | Aminotransferase, class I and II |
| Avin 15430 | 1515400 | 1515413 | 7 | - | 9.28571 | 8.68E-05 | 1 | GTCAGTAAATCGAA | | hypothetical protein |
| Avin 15440 | 1515400 | 1515413 | 254 | - | 9.28571 | 8.68E-05 | 1 | GTCAGTAAATCGAA | | methionine sulfoxide reductase B |
| Avin 27150 | 2785625 | 2785638 | 237 | - | 9.28571 | 8.68E-05 | 1 | GTCAGTAAATCGAA | | carboxymuconolactone decarboxylase-like protein |
| Avin 18470 | 1828210 | 1828223 | 377 | + | 9.27551 | 8.73E-05 | 1 | GTTAATAGTGTCAC | | histone-like protein |
| Avin 33210 | 3395980 | 3395993 | 46 | + | 9.26531 | 8.76E-05 | 1 | GTCGATTGTTCGTT | *aroF* | phospho-2-dehydro-3-deoxyheptonate aldolase |
| Avin 33220 | 3395980 | 3395993 | 44 | + | 9.26531 | 8.76E-05 | 1 | GTCGATTGTTCGTT | | hypothetical protein |
| Avin 10610 | 1007035 | 1007048 | 192 | - | 9.2551 | 8.78E-05 | 1 | GTCTCTATATCGGC | *tonB* | TonB protein |
| Avin 13440 | 1310352 | 1310365 | 169 | + | 9.2551 | 8.78E-05 | 1 | GCCAGATATTCGGC | *pdxH* | pyridoxamine 5-phosphate oxidase |
| Avin 47760 | 4845457 | 4845470 | 76 | + | 9.22449 | 8.91E-05 | 1 | GTTGCAAATCTGAT | | Conserved hypothetical protein |
| Avin 51720 | 5268477 | 5268490 | 116 | - | 9.22449 | 8.91E-05 | 1 | GCCAGAAAAGTCGC | *recC* | exodeoxyribonuclease V, gamma subunit |
| Avin 39540 | 4003887 | 4003900 | 171 | - | 9.18367 | 9.03E-05 | 1 | GTCGCTTGTTCGGC | *trmD* | tRNA (guanine-N(1)-)-methyltransferase |
| Avin 07210 | 684124 | 684137 | 381 | + | 9.16327 | 9.10E-05 | 1 | GTCATAAAAGTGCC | | hypothetical protein |
| Avin 07220 | 684124 | 684137 | 118 | + | 9.16327 | 9.10E-05 | 1 | GTCATAAAAGTGCC | | hypothetical protein |
| Avin 02800 | 265147 | 265160 | 345 | - | 9.15306 | 9.15E-05 | 1 | GCCGCAATATCGGC | | Endoribonuclease L-PSP family protein |
| Avin 32910 | 3371263 | 3371276 | 375 | + | 9.15306 | 9.15E-05 | 1 | GCCGATAGATCCGC | | Ribonuclease BN protein |
| Avin 07220 | 684174 | 684187 | 68 | + | 9.10204 | 9.30E-05 | 1 | GCCATTATTTTGGC | | hypothetical protein |
| Avin 31780 | 3287325 | 3287338 | 83 | + | 9.10204 | 9.30E-05 | 1 | GTCAAAAATTCCGT | | hypothetical protein |
| Avin 31790 | 3287325 | 3287338 | 130 | + | 9.10204 | 9.30E-05 | 1 | GTCAAAAATTCCGT | | hypothetical protein |
| Avin 13820 | 1343015 | 1343028 | 265 | - | 9.08163 | 9.39E-05 | 1 | GTCACCAGAGTGAC | *recO* | DNA repair protein RecO |
| Avin 41760 | 4208971 | 4208984 | 257 | + | 9.08163 | 9.39E-05 | 1 | GCCAACAGATCGAC | *adh* | 2,3-butanediol dehydrogenase |
| Avin 12400 | 1205516 | 1205529 | 22 | + | 9.07143 | 9.43E-05 | 1 | GTCGGAATTCCGGC | | PepSY-associated TM helix |
| Avin 10880 | 1037836 | 1037849 | 110 | + | 9.06122 | 9.47E-05 | 1 | GCCGGTATATCGAT | *algV* | Alginate biosynthesis protein AlgV |
| Avin 13960 | 1363550 | 1363563 | 41 | - | 9.06122 | 9.47E-05 | 1 | GCTAATTATCCGAC | | Glycosyl transferase, family 51 |
| Avin 12010 | 1157064 | 1157077 | 329 | - | 8.96939 | 9.79E-05 | 1 | GCCGGATGATCGGC | | conserved hypothetical protein |
| Avin 12020 | 1157064 | 1157077 | -37 | - | 8.96939 | 9.79E-05 | 1 | GCCGGATGATCGGC | | N-acetyltransferase (GNAT) family |
| Avin 12030 | 1157064 | 1157077 | 385 | - | 8.96939 | 9.79E-05 | 1 | GCCGGATGATCGGC | | conserved hypothetical protein |
| Avin 20570 | 2047912 | 2047925 | 160 | - | 8.94898 | 9.87E-05 | 1 | GCTGATAAAGTGGC | | VacJ-like lipoprotein |
| Avin 17910 | 1780597 | 1780610 | 305 | - | 8.93878 | 9.90E-05 | 1 | GTTGGAAAAGCGGC | | Glycosyl transferase, group 1 family protein |
| Avin 22590 | 2255867 | 2255880 | 109 | - | 8.93878 | 9.90E-05 | 1 | GCCGGTAGTCCGGC | | Molybdenum-pterin binding domain protein |
| Avin 52190 | 5334153 | 5334166 | 26 | + | 8.91837 | 9.98E-05 | 1 | GCCAGTTTTTCGAC | *atpH* | F1 sector of membrane-bound ATP synthase, delta subunit |
| Avin 06940 | 659684 | 659697 | -21 | - | 8.90816 | 0.0001 | 1 | GCTGCAAATCCGAC | *prmA* | ribosomal protein L11 methyltransferase |

***The start and end coordinates refers to the genome sequence of *A. vinelandii* DJ strain (Genbank accesion: CP001157)**

*P. aeruginosa* FleQ logo

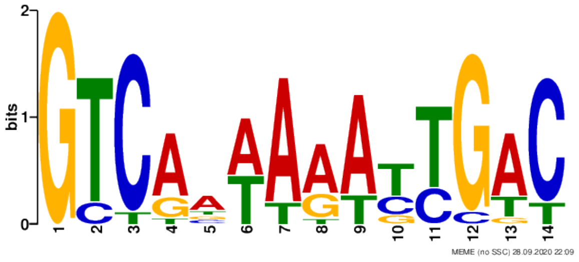
bits
MEME (no SSC) 28.09.2020 22:09

gase
